# Spatial cancer systems biology resolves heterotypic interactions and identifies disruption of spatial hierarchy as a pathological driver event

**DOI:** 10.1101/2023.03.01.530706

**Authors:** Fabian V. Filipp

**Affiliations:** Cancer Systems Biology, Institute of Diabetes and Cancer, Helmholtz Zentrum München, Ingolstädter Landstraße 1, D-85764 München, Germany; School of Life Sciences Weihenstephan, Technical University München, Maximus-von-Imhof-Forum 3, D-85354 Freising, Germany; Institute for Advanced Study, Technical University München, Maximus-von-Imhof-Forum 3, D-85354 Freising, Germany; Metaflux, San Diego, CA, 92105, United States of America

**Keywords:** spatial transcriptomics, single-cell transcriptomics, spatial, single-cell, multi omics, systems biology, precision medicine, cell communication, epithelial‐to‐mesenchymal transition, tumor immune microenvironment, melanoma, cancer, skin, dermatology, skcm, frizzled, wnt, vangle, catenin, non-canonical, spRNA-Seq, scRNA-Seq, IO, SC, AI, ML, DL, DNN, PCP, EMT, ECM, TME, TIME, TCGA, SKCM, FZD, DVL, VNGL, CTNNB1

## Abstract

Spatially annotated single-cell datasets provide unprecedented opportunities to dissect cell-cell communication in development and disease. Heterotypic signaling includes interactions between different cell types and is well established in tissue development and spatial organization. Epithelial organization requires several different programs that are tightly regulated. Planar cell polarity (PCP) is the organization of epithelial cells along the planar axis, orthogonal to the apical-basal axis. Here, we investigate PCP factors and explore the implications of developmental regulators as malignant drivers. Utilizing cancer systems biology analysis, we derive a gene expression network for WNT-ligands (WNT) and their cognate frizzled (FZD) receptors in skin cutaneous melanoma. The profiles supported by unsupervised clustering of multiple-sequence alignments identify ligand-independent signaling and implications for metastatic progression based on the underpinning developmental spatial program. Omics studies and spatial biology connect developmental programs with oncological events and explain key spatial features of metastatic aggressiveness. Dysregulation of prominent PCP factors such as specific representatives of the WNT and FZD families in malignant melanoma recapitulates the development program of normal melanocytes but in an uncontrolled and disorganized fashion.

## Introduction

### Spatial single-cell transcriptomics sheds light on intercellular crosstalk

The emerging fields of single-cell RNA-sequencing (scRNA-Seq) and spatial transcriptomics (spRNA-Seq or spRNA-FISH, for spatial whole transcriptome or spatial fluorescence in situ sequencing imaging approaches, respectively) combine the benefits of traditional histopathology with single-cell gene expression profiling. The ability to connect the spatial organization of molecules in cells and tissues with their gene expression state enables the mapping of developmental stages as well as resolving specific disease pathologies. spRNA-Seq has the ability to decode molecular proximities from sequencing information and construct images of gene transcripts at sub-cellular resolution. As a result, tissue heterogeneity and intercellular crosstalk can now be charted and delineated with never-before-seen accuracy.

### Tissue development relies on tightly controlled spatial programs

Spatially annotated single-cell datasets, omics-enhanced imaging data coupled with spRNA-Seq and/or sorted scRNA-Seq, provide unprecedented opportunities to dissect cell-cell communication. In skin biology as well as in clinical dermatology, this is particularly useful to advance our understanding of the epithelial tissue program in dermal development and disease (Cang, 2023). Epithelial organization requires several different programs that are tightly regulated. Planar cell polarity, PCP, is the organization of epithelial cells along the planar axis orthogonal to the apical-basal axis. Planar cell polarity is observed in an array of developmental processes that involve collective cell movement and tissue organization, and its disruption can lead to severe developmental defects. Recent research on flies and vertebrates has discovered new functions for planar cell polarity, as well as new signaling components and mechanistic models. However, despite such progress, the search to simplify principles of understanding continues, and important mechanistic uncertainties still pose formidable challenges. Cell migration is a highly integrated multistep process that orchestrates embryonic morphogenesis, contributes to tissue repair and regeneration, and drives disease progression in cancer, mental retardation, atherosclerosis, and arthritis. The migrating cell is highly polarized with complex regulatory pathways that spatially and temporally integrate its component processes. Neural crest cells are a well-organized, pluripotent, yet temporary group of cells from the embryonic ectoderm that have an ability to migrate and give rise to diverse cell lineages, including neurons, Schwann cells, and melanocytes (Biermann, 2022). The neuroectodermal developmental program is attributed to the elevated propensity of melanoma cells for organotropism and brain metastases.

### Loss of differentiation and spatial organization in cancer

Tissue homeostasis is supported by cellular growth and proliferation. Size is a fundamental attribute impacting molecular structure, cellular design, fitness, and function. Concentration-dependent cell size check points ensure that cell division is delayed until a critical target size has been achieved. Likewise, control of cell proliferation is a fundamental aspect of tissue formation in development and regeneration. Stem cells exhibit a low proliferation rate while maintaining a high proliferative capacity and are often small. In contrast, cancer cells do not precisely reverse this process yet show characteristics of loss of differentiation. Extracellular signals or ligand-receptor interactions may independently induce growth and division. Specific driver genes have been linked to specific aspects of differentiation loss and spatial organization (Hodis, 2021).

### Delineation of molecular driver events by understanding the molecular hierarchy of cancer

Melanoma is an aggressive type of skin cancer that partially recapitulates the development program of normal melanocytes but in an uncontrolled and disorganized fashion. Tissue disorganization is one of the main hallmarks of cancer. Polarity proteins are responsible for the arrangement of cells within epithelial tissues through the asymmetric organization of cellular components. Consistent with these findings are recent studies investigating control elements of melanocyte and skin tissue formation (Li, 2019; Dong, 2023). If we understand the cellular hierarchy in melanoma, wherein growth and metastasis are governed by the rules of the developing embryonic neural crest, we can delineate oncological drivers disrupting tissue homeostasis. Spatial single-cell profiling has been invaluable in quantifying the unprecedented heterogeneity and plasticity of malignant melanoma. A systems biology analysis revealed distinct subpopulations representing lineage-specific melanocyte inducing transcription factor (MITF) as well as mesenchymal-like clones with high proliferative potential (Karras, 2022; Biermann, 2022). Taken together, signaling and transcriptional elements govern a molecular hierarchy in melanoma that uncouples growth and metastasis.

## Results

### Recapitulation of a developmental program in cancer metastasis

Early on, in the history of science, the fields of evolutionary developmental biology and oncology shook hands when they created the portmanteau WNT from the planar cell polarity drosophila wingless, WG, mutant and the mouse mammary tumor virus integration, INT, site. WNT-ligands are recognized by cognate frizzled (FZD) receptors. Today, the WNT, FZD, and beta-catenin signaling pathways constitute an evolutionarily conserved cell-cell communication system that is important for stem cell renewal, cell proliferation, and cell differentiation both during embryogenesis and during adult tissue homeostasis (Figure 1). The reliance on specific protein isoforms highlights a cellular ability to fine-tune this signaling pathway. On the one hand, WNT- and FZD-dependent interactions may lead to cytoplasmic stabilization of soluble β-catenin, classified as the canonical WNT pathway. On the other hand, receptor activation may result in Planar cell polarity or calcium activation, both of which classify as non-canonical WNT pathways. Prominent yet highly selective dermal expression patterns warrant that in skin development, both canonical and non-canonical pathways play essential roles. The planar cell polarity arm of WNT signaling is critical during melanocyte specification from the neural crest. Not surprisingly, the cellular expression pattern within skin cancer cells as well as surrounding microenvironment components recapitulates such distinct development programs hijacking cellular tracks of tissue remodeling and facilitating mobility and metastatic transition.

**Figure 1:**
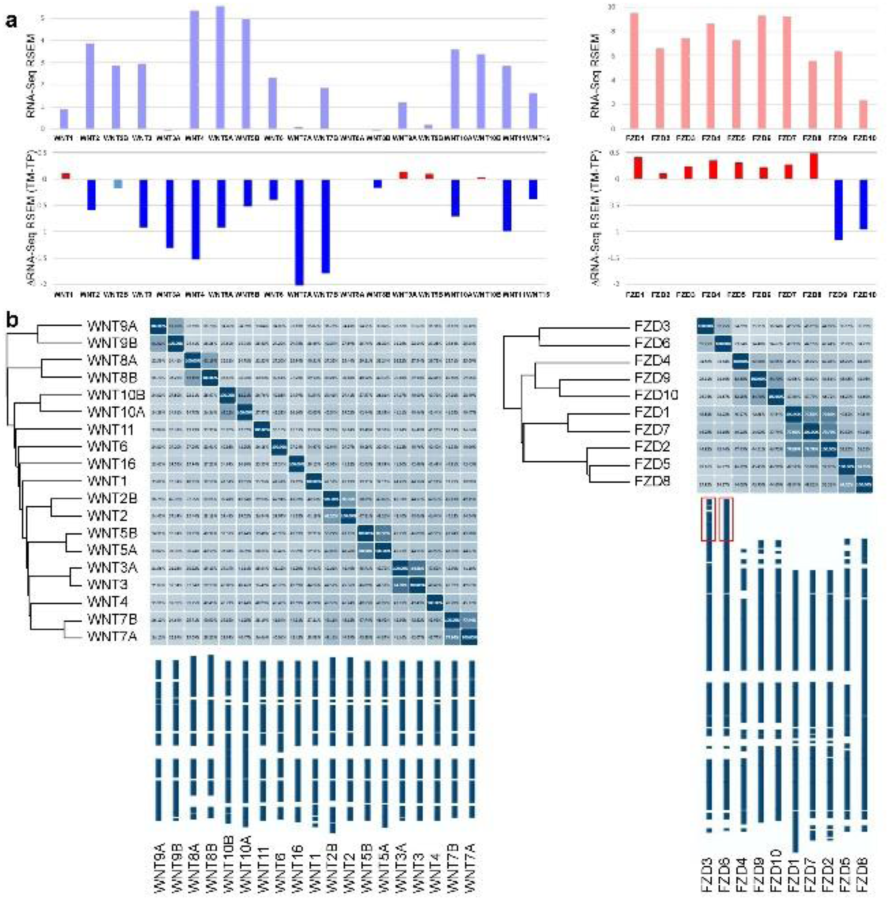
WNT-ligand (WNT) and cognate frizzled (FZD) receptor gene expression network in skin cutaneous melanoma Disruption of heterotypic signaling in skin cutaneous melanoma

Assisted by structure-based modeling and comparative multiple sequence alignments, cancer system biology can break down the highly conserved WNT molecular network, which encompasses 19 closely related WNT ligands (WNT1–16, including A and B isoforms) and 10 Frizzled receptors (FZD1–10) that direct the self-renewal and regeneration of many tissues during their development. FZDs are G protein-coupled receptors characterized by seven transmembrane-spanning domains, a cysteine-rich N-terminal ligand binding domain, and a C-terminal intracellular activation domain. Depending on their activation, ligands, and intracellular binding adapter proteins, FZDs are capable of transmitting extracellular signals into diverse transcriptional program outputs that determine cell fate during normal and pathogenic development. A common feature of the WNT-FZD gene expression network in cancer is upregulation and hyperactivation of both classes of oncogenes. Heterotypic interactions strictly rely on the juxtaposed expression of receptor and ligand on different cell types. In contrast, cancer cells express both molecules, even several isoforms of receptor and ligand, at the same time, overcoming any spatial constraints of tissue regulation.

## Methods

### A finely tuned gene expression network of ligand and cognate receptor isoforms is dysregulated in skin cutaneous melanoma

RNA-Seq gene expression data from 302 patient specimens, including normal, tumor, and metastatic tissue, was subjected to differential gene expression analysis. Read counts were scaled via the median of the geometric means of fragment counts across all libraries. Transcript abundance was quantified using normalized single-end RNA-Seq reads in read counts as well as reads per kilobase million (RPKM). Since single-end reads were acquired in the sequencing protocol, quantification of reads or fragments yielded similar results. Statistical testing for differential expression was based on read counts and performed using EdgeR in the Bioconductor toolbox. Gene families were further analyzed using the EMBL phylogenetics Smart Domain database and Clustal X 2.0, a multiple sequence alignment program that takes advantage of guide trees and Hidden Markow Model profile-profile techniques to generate alignments between sequence families. The study was carried out as part of an IRB-approved study, dbGap ID 5094 “Somatic mutations in melanoma” (Guan, 2015). The results shown are in part based upon data generated by The Cancer Genome Atlas (TCGA) Research Network, http://cancergenome.nih.gov. Restricted-access whole-genome sequences and whole-exome sequences were obtained from the TCGA data portal.

## Discussion

### Planar cell polarity regulators with distinct roles in cancer metastasis

*FZD3* and *FZD6* share high sequence homology and function through the noncanonical WNT pathway to play a critical role in migration and planar pattern formation. Structure-based sequence analysis identifies a distinct 100 amino acid C-terminal extension, exclusively found in FZD3 and FZD6 (Figure 1). This extra tail makes FZD3 and FZD6 proteins special among their peers, enabling unique cell-cell signaling properties. Cancer genomics leverages a genome-wide insight and reveals that dysregulation of the WNT-FZD oncoprotein network is a driver of distinct spatial, developmental, and pathological programs.

### Implications for epithelial-to-mesenchymal transition and the tumor immune microenvironment

A *FZD3* knockout revealed the underlying transcriptional regulators of the *TWIST* and *SNAIL* families (Li, 2019). *TWIST* and *SNAIL* are commonly defined not only as developmental markers of neural crest cells in dorsolateral migration but also as drivers of proliferation and invasion during malignant transformation (Figure 2). Dysregulation of *SNAIL* transcription factors upon *FZD3* knockout is common to the *FZD6* knockout model. In contrast, downregulation of epithelial-to-mesenchymal transition-related *ZEB* homeobox transcription factors is specific to the *FZD6* knockout model. Strikingly, knockout of *FZD6* did not affect primary tumor formation or cellular proliferation but inhibited distant metastasis and cellular migration of melanoma into distant tissues (Dong, 2023). Because of similar pivotal roles in tissue polarity, sequence homology, and overexpression in melanoma, it may not come as a surprise that *FZD3* and *FZD6* display some expected functional redundancy. However, despite such similarities, the oncogenes *FZD3* and *FZD6* play quite different molecular roles in melanoma progression. *FZD3* promotes melanoma metastasis by stimulating cell cycle progression, while *FZD6* regulates cell invasiveness (Figure 2).

**Figure 2:**
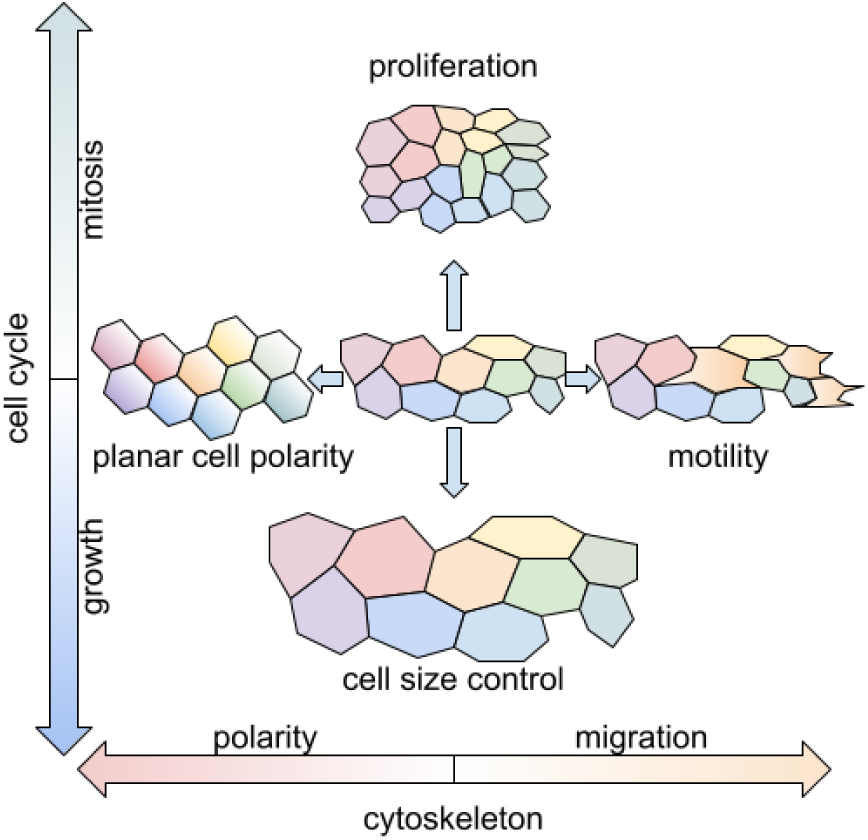
Spatial single-cell profiling identifies disruption of cellular hierarchy in cancer

### Omics-based spatial system biology brings tissue control elements and metastasis drivers to the surface

In cancer, the FZD3 receptor oncoprotein signals independently of the canonical WNT pathway, while FZD6 may also engage canonical WNT signaling. The data identifies significant functional specificity of spatial regulators in cancer, despite similar roles in cell planarity and tissue development (Figure 2). With the advent of spatial omics, it is now possible to link latent developmental programs with oncological events and explain key spatial features of metastatic aggressiveness.

## Conclusions

### Spatial omics to overcome the developmental immune privilege in disease

Facilitated by spatial omics insight, the developmental path might also have important implications for stem cell biology and immuno-oncology. Tissue aging, cellular senescence, and controlled cell death have an underappreciated immunological control element. Similarly, evasion of cancer cells from the endogenous immune repertoire can be attributed to an immune privilege that only sheltered stem cell populations enjoy, such as migrating precursors, but foreign cells abuse as a hard-to-overcome escape pathway. T lymphocytes are the primary effectors of the anti-tumor response, but the interplay between melanoma and the immune system is complex, dynamic, and incompletely understood. Sustained progress in unraveling the pathogenesis of melanoma regression has led to the identification of therapeutic targets, culminating in the development of immune checkpoint inhibitors for the management of advanced disease.

### Spatial regulators as predictive biomarkers and therapeutic targets

The first anti-FZD antibodies or engineered FZD-Fc fusion proteins, which are serving as decoy receptors for WNT ligands, are in clinical trials for patients with advanced solid tumors. Modern techniques allow for high-resolution spatial analyses of the tumor immune microenvironment (Filipp, 2019; Reynolds, 2021; Schäbitz, 2022). Such spatial omics studies may lead to a better understanding of the immune drivers of melanoma regression. As a result, they aid in the search for new prognostic and predictive biomarkers in the treatment of migratory metastatic cell populations (Hodis, 2022; Cang, 2023). Going forward, spatial omics will guide clinical decision-making to create durable anti-cancer therapy responses.

## Abbreviations

FZD: frizzled receptor
WNT: wingless-related integration site ligand (portmanteau of wg and int)
PCP: planar cell polarity
EMT: epithelial‐to‐mesenchymal transition
ECM: extracellular matrix
TME: tumor microenvironment
TIME: tumor immune microenvironment
scRNA-Seq: single-cell RNA-sequencing
spRNA-Seq: spatial transcriptomics

## Declarations

### Data availability statement

The manuscript was made publicly available to the scientific community on preprint servers bioRxiv at https://doi.org/10.1101/2023.03.01.530706 and arXiv at https://doi.org/10.48550/arXiv.2303.00933.

